# A panel of Diversity Outbred mouse embryonic stem cells: enabling the functional characterization of natural genetic variation *in vitro*

**DOI:** 10.64898/2026.09.11.747888

**Authors:** Morgan M. Brady, Emily Swanzey, Kathryn Janeczko, Samuel J. Widmayer, Hannah B. Dewey, Selcan Aydin, Whitney Martin, Catherine Brunton, Anuj Srivastava, Daniel M. Gatti, Laura G. Reinholdt, Steven C. Munger, Christopher L. Baker

## Abstract

Phenotypic variation in pluripotent stem cells arises largely from genetic variants that impact stability of the pluripotent state, response to differentiation cues, and cell fate diversification. Panels derived from genetic reference populations leverage genetic diversity to map the loci influencing these molecular and cellular traits. Here we describe the first publicly available resource for *in vitro* systems genetics in *Mus musculus*: 299 genetically diverse, highly heterozygous embryonic stem cell lines derived from Diversity Outbred mice, with minimal kinship across the panel. We characterize genomic integrity and allelic variation across the panel and describe differences in pluripotency and differentiation networks in embryoid bodies. Mapping the genetic determinants of gene expression differences in response to cadmium exposure in embryoid bodies identified 13 regulatory hotspots, the largest acting on ribosome biogenesis genes. This panel enables mapping of loci and gene-by-environment interactions that influence molecular identity in pluripotent cells and their derived cell types.

**Highlights:**

- First publicly available resource for *in vitro* system genetics in *Mus musculus*
- 299 Diversity Outbred-derived mESCs recapitulate allelic diversity, low kinship
- Gene expression analysis shows variation in pluripotency, differentiation networks
- Panel enables high-resolution mapping of loci and G×E interactions

**eTOC:** Systems genetics resources are needed to identify genes and regulatory networks contributing to complex traits across genetically diverse populations. Baker and colleagues generated, genotyped, and characterized a panel of Diversity Outbred-derived mESCs that maintains the genetic heterozygosity of the source population for use in high-resolution mapping of loci responsible for developmental and G×E phenotypes, enabling *in vivo* functional validation.

## Introduction

Embryonic stem cells can differentiate into a wide range of biomedically relevant cell types and organoid cultures, making them a foundational tool for regenerative medicine and disease modeling. Much of this work, however, relies on cells from a limited number of genetic backgrounds, obscuring how genetic variation between individuals shapes responses to differentiation cues and other cellular behaviors. Panels of genetically diverse stem cell lines address this directly, enabling screens for variable responses to perturbation and discovery of loci that modify cell state and differentiation outcomes. Realizing this potential has followed from advances in functional genomic technologies, including high-throughput sequencing and mass spectrometry, which together enable extensive molecular trait capture and quantitation across many lines. Integrating these datasets using systems genetics leverages genetic variation to resolve how molecular networks shape organism-level traits and, ultimately, contribute to complex human diseases. Accordingly, recent work has identified loci influencing variation in pluripotency and cell state transition in both human^1,2^ and mouse^3–6^ cell models, including loci that regulate many genes at once.

The power of a mapping panel, however, depends on how it is constructed. Many variants only subtly influence a trait of interest and necessitate the use of large panels to detect small effects in aggregate, while rare variants with strong effect sizes may be underrepresented and therefore difficult to detect^7^. Relatedness among lines imposes a further limit, as lines sharing long haplotypes contribute redundant information and reduce both power and the ability to resolve a locus. Due to historical reliance on isogenic strains, model organism studies are often limited in genetic variability, making generalization of their results to genetically diverse populations a challenge^8^. The Diversity Outbred (DO) population was designed to address these limitations. Maintained as a randomized outbred stock descended from eight inbred founder strains, each DO animal carries a unique and highly heterozygous genome^9^. Randomized outcrossing maintains balanced founder allele frequencies, increasing the prevalence of rare variants while successive generations narrow haplotypes through accumulated recombination. Because all eight founder genomes are sequenced, haplotype effects estimated at a locus can be resolved to individual variants. These properties have supported mapping of traits spanning physiology^10–12^, behavior^13–15^, disease susceptibility^16–18^, lifespan^19,20^, regenerative biology^21^, and exposure response^22,23^.

More recently, studies utilizing DO-derived cell types examined LIF response in mESCs^4^, toxicant response in fibroblasts^24^, and bone mineral density in stromal cells^25^. To extend this approach to a well-characterized, publicly available resource, we derived a large DO mouse embryonic stem cell (DO mESC) panel, comprised of 299 sex-balanced lines, each established from a unique, non-sibling male/female DO mating to minimize kinship and maximize genetic diversity. Each DO mouse is a genetically unique individual whose combination of alleles cannot be recovered once the animal is gone; pluripotent cell lines capture this diversity into a renewable resource, such that once genotyping is performed on the cell line, the experimental burden is reduced and multiple phenotypes can be compared across studies. Further, all DO mESCs can be grown in defined feeder-free, serum-free culture conditions, limiting experimental variability. Human iPSC collections provide genetically diverse reference lines as a similar resource^26,27^, but donors carry alleles at their frequency in the source population, limiting mapping power to detect rare variants. Because founder allele frequencies in the DO are balanced by design, the DO mESC panel is therefore complementary, and loci identified in the panel can be further tested *in vivo*^7,17,28^.

Here we describe the systematic generation and characterization of this DO mESC resource. Panel lines were genotyped using the Giga Mouse Universal Genotyping Array (GigaMUGA)^29^, and the resulting genotypes are available for mapping any trait measured in these lines. Following differentiation to embryoid bodies (EBs), gene expression analysis revealed substantial transcriptional heterogeneity between lines in the differentiated state that is not present in the naïve state. Finally, to demonstrate the utility of this panel for perturbation screening, we differentiated 200 DO mESC lines into EBs in the presence or absence of cadmium (Cd), a human carcinogen and one of the top 10 chemicals of public health concern identified by the World Health Organization^30^. We performed gene-by-environment (G×E) QTL mapping and identified 13 genomic hotspots where genetic variation shapes gene expression response to cadmium during early cell state transitions.

## Results

### Panel design and generation

The Diversity Outbred (DO) mouse population is a highly intercrossed heterogeneous stock derived from the same eight inbred founder strains used to create the Collaborative Cross (CC) recombinant inbred strains. These include five common laboratory strains (A/J, C57BL/6J, 129S1/SvImJ, NOD/ShiLtJ, and NZO/HlLtJ) and three wild derived inbred strains (CAST/EiJ, PWK/PhJ, and WSB/EiJ) that together segregate nearly 50 million single nucleotide polymorphisms (SNPs) and capture nearly 90% of known genomic variation in laboratory mice^9^. The DO population was generated by random outcrossing of partially inbred “pre-CC” mice to create mice with broadly distributed genetic diversity, a high level of allelic heterozygosity, and balanced founder allele frequencies, with frequencies >5% for most SNP minor alleles (Figure 1A). The DO mice used here for ESC derivation were outcrossed at generations 40–50, yielding cell lines that have an accumulated 997 meiotic recombination events on average (range 809–1210 detected crossovers). Importantly, the low linkage disequilibrium in these highly recombined mESCs make them an ideal resource for high resolution genetic mapping studies.

**Figure 1.**
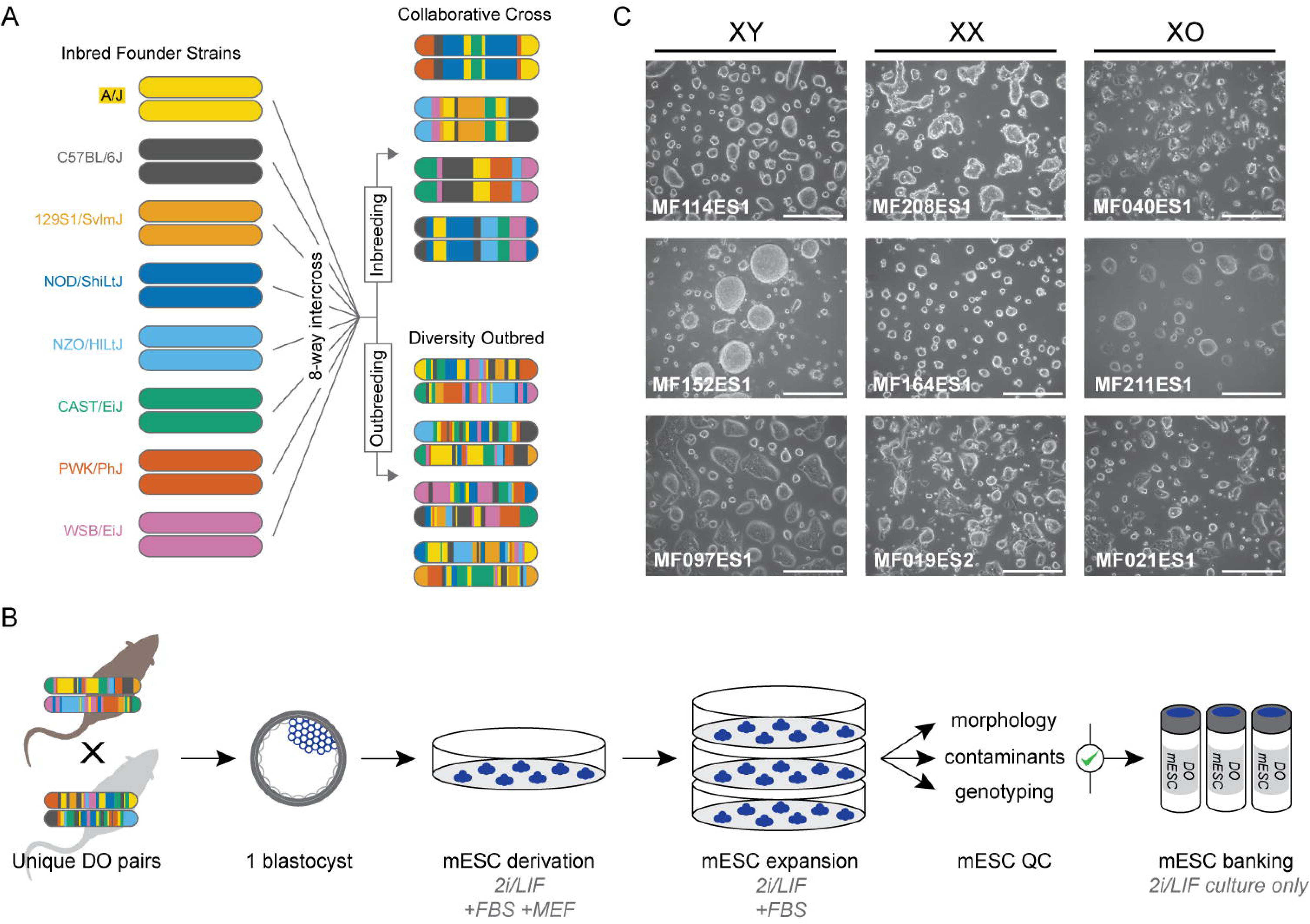
Derivation of the DO mESC resource. (A) Overview of the DO breeding scheme. Founder strains are listed on the left. DO mice were originally derived from incipient Collaborative Cross (CC) strains and are maintained as heterogenous stocks through a randomized breeding scheme. (B) Overview of the DO mESC derivation protocol. Each DO mESC line was derived from unique, non-sibling matings, then expanded and tested for contaminants before GigaMUGA genotyping and cryovial banking. (C) Diverse DO mESC morphology from selected lines at P6. Scale bar set to 500 µm. Despite differences in colony size and shape, most lines display the typical discrete, refractile ESC colony morphology indicative of pluripotency.

To create the DO mESC panel, we crossed unique, non-sibling male/female DO pairs and derived one primary cell line per mating, thereby maximizing genetic diversity and reducing kinship (Figure 1B). Two blastocysts were collected from each mating at embryonic day 3.5 (E3.5) using previously described protocols^31^ and modified media formulations, with most lines successfully derived from the first embryo selected. A small number of primary line derivations (*n* = 16) were initially unsuccessful and required re- derivation from the secondary, banked “sibling” blastocyst. A total of 299 DO mESC lines were banked once cultures were established (P6–P12; >90% P6–P9) and tested negative for bacterial, fungal, and mycoplasma contaminants (Supplemental Table 1). All panel lines can be grown in identical and defined feeder-free, serum-free 2i/LIF culture conditions.

Consistent with findings from previous work^3–5^, DO mESC lines displayed variability in colony morphology and growth rate (Figure 1C), likely driven by their diverse genetic background and indicative that their phenotypic differences can be mapped back to the DO founder population. Most lines were characterized by typical refractile, droplet-like colonies with smooth or jagged edges, while a smaller number of colonies displayed flattening peripheral cells, demonstrating the wide variety of morphological phenotypes that are observable and well-tolerated in defined, controlled culture conditions.

### Assessing panel relatedness

To genotype and estimate kinship for DO mESC lines, we utilized the GigaMUGA genotyping platform. This high-density array comprises 141,090 SNP markers that are evenly distributed across the mouse genome and were selected to be informative for inferring founder strain ancestry in the DO population. We observed minimal autosomal kinship between DO mESC lines across the entire panel—underscoring a key advantage of our mating scheme that employed unique DO parents for each mESC line (Figure 2A). Indeed, the observed kinship in this panel is far lower than previous reported DO mESC panels derived from fewer DO founders and earlier generations^4,32^. Next, we determined the sex chromosome content of each line using GigaMUGA probe intensity of X- and Y-linked markers, identifying 162 XY, 131 XX, 1 potentially XXY, and 5 XO/ambiguous lines (Figure 2B, Table 1). The latter is indicative that a subset of lines spontaneously lost an X chromosome during the derivation or culture process (Figure 2B), a commonly reported aneuploidy in mESC culture due to the instability of the XX karyotype^33,34^.

**Figure 2.**
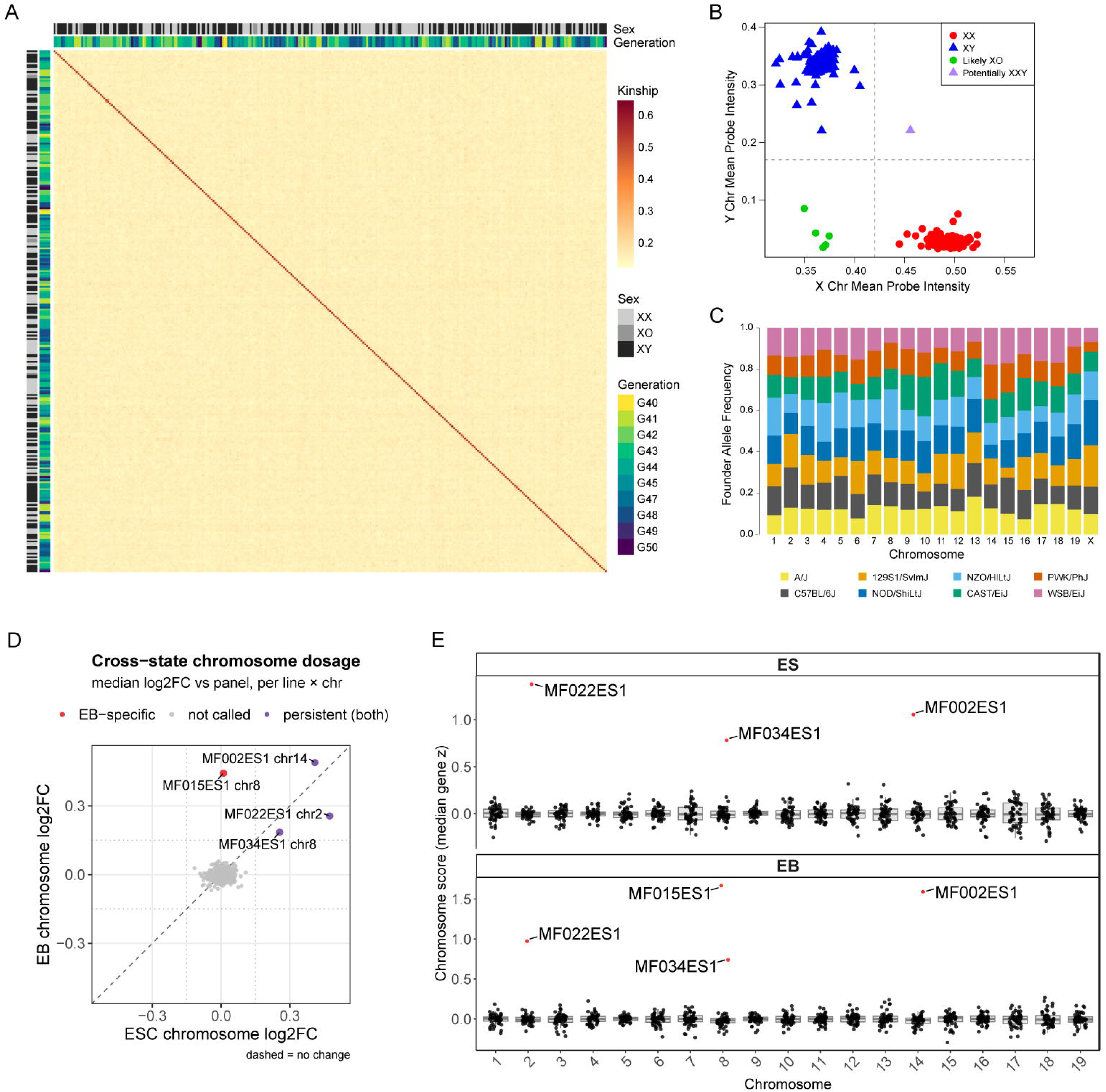
Assessing DO mESC panel relatedness and genomic quality. (A) GigaMUGA autosomal SNP genotypes show minimal kinship across the 299 DO mESC panel, with no kinship structure associated with outbreeding generation or chromosomal sex observed. Pair-wise kinship values derived from correlation of autosomal SNPs are plotted as a heatmap, and mESC chromosomal sex and outbreeding generation are indicated above and beside the heatmap. (B) Mean GigaMUGA probe intensity of X/Y-linked markers show panel lines are sex-balanced, with 162 XY, 131 XX, 5 XO/ambiguous lines, and 1 potential XXY line. (C) Calculated genome-wide DO founder allele frequencies found small but significant deviations from the expected 0.125 frequency of PWK/PhJ (0.108) and NOD/ShiLtJ (0.14). When partitioned by chromosome, slight significant deviations were found on Chrs 2, 5–6, 8, 10–11, 13–15, and X (Chi-squared, BH adjusted p-value < 0.05). (D) Gene expression data from 48 DO mESC and corresponding differentiated EB samples found 4/48 lines harbored autosomal aneuploidies on Chrs 2, 8, and 14 across states (purple) or in EB samples only (red) (|robust z| ≥ 4 and |log2FC| ≥ 0.15). Each point represents one line per autosome; the x-axis is ESCs and y-axis is EBs for that same line and chromosome (48 lines × 19 autosomes = 912 points). Points near the dashed diagonal have similar values between cell states. (E) Per-chromosome distributions of the chromosome score in each state. The values for most chromosomes form a tight cloud near zero; flagged aneuploidy calls are labeled.

**Table 1.**
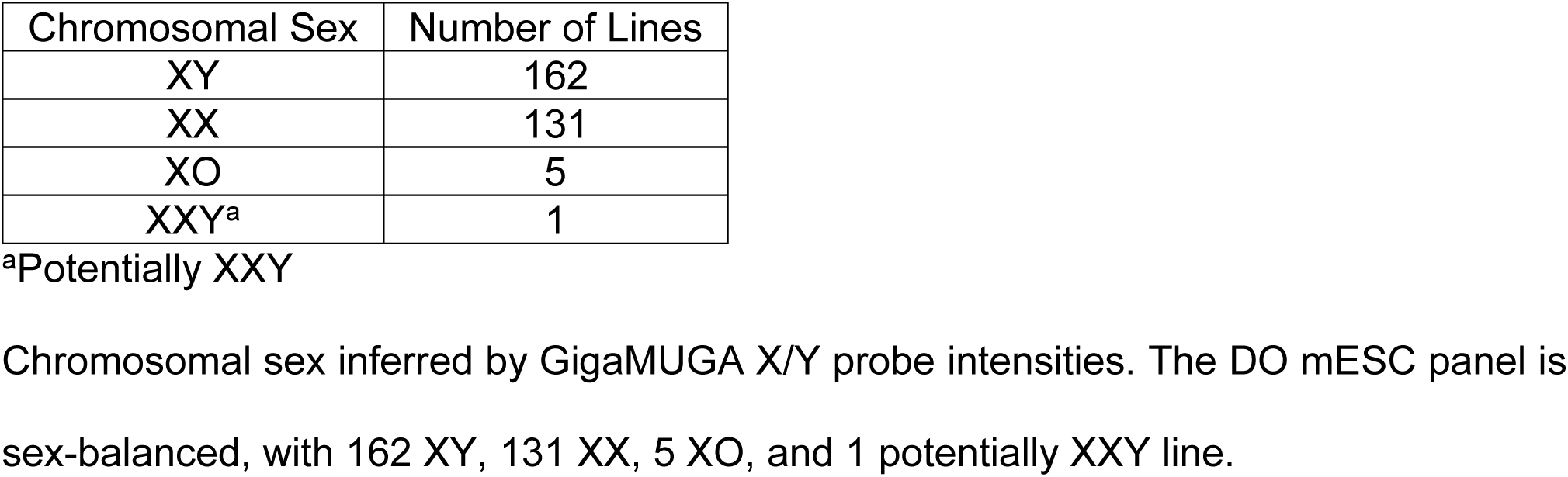
Chromosomal sex of DO mESC panel lines.

| Chromosomal Sex | Number of Lines |
| --- | --- |
| XY | 162 |
| XX | 131 |
| XO | 5 |
| XXY<sup>a</sup> | 1 |

<sup>a</sup>Potentially XXY
Chromosomal sex inferred by GigaMUGA X/Y probe intensities. The DO mESC panel is sex-balanced, with 162 XY, 131 XX, 5 XO, and 1 potentially XXY line.

Further, we assessed the relative balance of founder alleles present in the DO mESC lines. The large breeding population and random outbreeding scheme ensures that founder allele frequencies are largely balanced across the population, with a minor allele frequency (MAF) ≥ 1/8^35^. We calculated founder allele frequency genome-wide and found small but statistically significant deviations from the expected 0.125, driven mostly by under-representation of PWK/PhJ (0.108) and over-representation of NOD/ShiLtJ (0.14). A similar allele frequency is observed in a recently derived DO iPSC panel reprogrammed from tail-tip fibroblasts at similar outbred generations^36^, and likely reflects founder allele deviations present in the breeding population rather than allelic incompatibilities in the derivation of mESCs. Partitioning founder allele frequencies by chromosome reveals small but significant deviations from expectation on chromosomes 2, 5–6, 8, 10–11, 13–15, and X (Chi-squared, BH adjusted p-value < 0.05; Figure 2C).

### Aneuploidy screening

Embryonic stem cells can acquire chromosomal abnormalities during *in vitro* culture, due to serial passaging or prolonged *Mek1/2* suppression^37,38^. mESC chromosome copy number has previously been linked to rapid growth that can confer selective advantages in cell culture, particularly trisomy 8 and 11^39–41^. Additionally, aneuploid mESC lines are associated with low germline transmission^39^, differentiation defects^40,41^, and sex-linked dosage modulation^34^. Controlling chromosome copy numbers is therefore critical for efficacy, reproducibility, and to ensure observed effects of interest are caused by genetic background.

To determine if DO mESC lines carried whole-chromosome aneuploidies and whether these imbalances differ between cell states, we screened 48 naïve DO mESC lines and their matched EBs using gene expression data. Whole-chromosome gains and losses produce coordinated shifts in expression across all genes on the affected chromosome, allowing dosage imbalance to be inferred. For each gene, log-CPM values were centered to the panel median, and an autosome-level score was calculated as the median of these gene-level differences across all genes on the chromosome. Autosomes were flagged only when they showed both an extreme outlier score (|robust z| ≥ 4) and a chromosome-wide shift of at least 0.15 log_2_ (∼11%). Three lines carried trisomies in the ESC state (MF002ES1, XY, Chr 14; MF022ES1, XY, Chr 2; and MF034ES1, XY, Chr 8), each of which persisted in the corresponding EBs, while one additional trisomy was detected only in the EB state (MF015ES1, XY, Chr 8) (Figure 2D–E, Supplemental Figure 1A, Supplemental Table 2).

We spot-checked these expression-based calls against orthogonal methods. GigaMUGA B-allele frequency (BAF) and log R ratio (LRR) profiles confirmed the Chr14 gain in MF002ES1 and the Chr2 gain in MF022ES1 but detected no copy number changes in MF015ES1 or MF034ES1 (Supplemental Figure 1C). Cytogenetic karyotyping of 20 lines independently identified aneuploidy (<70% euploid metaphases^31^) in MF015ES1 and MF022ES1, but not MF002ES1 (MF034ES1 was not karyotyped) (Supplemental Figure 1B). Karyotyping also identified two lines that expression-based screening did not flag: MF005ES1 (XX, 35% euploid, chromosome unknown) and MF010ES1 (XX, 50% euploid, chromosome unknown). Further, karyotyping identified MF021ES1 and MF040ES1 as XO, consistent with our GigaMUGA X/Y probe intensity data (Supplemental Table 3). Because DNA for genotyping was collected at line expansion and RNA later in culture, discordant calls are consistent with aneuploidies arising during passaging. In summary, 4/48 lines screened carried autosomal aneuploidy across both cell states.

### Characterizing variation in pluripotency and differentiation propensity

Genetic background influences pluripotency maintenance and differentiation propensity in both human^1,2^ and mouse^3–6^ cell models. EBs are a commonly used *in vitro* differentiation model formed by aggregation of ESCs into an organoid resembling the early blastocyst, making them well suited to assess pluripotency and lineage commitment in both human and mouse pluripotent cells^42^. To understand the extent of pluripotency and differentiation network variability within these genetically diverse lines, we expanded a subset of 48 DO mESC lines for genome-wide gene expression comparison in undifferentiated/naïve (ESC) and spontaneously differentiated (EB) states (Figure 3A). Early passage DO mESC cells were induced to differentiate in serum-containing media supplemented with bFGF and grown for 4 days (equivalent to E7.5–8.0), with each line contributing up to 24 pooled EBs for RNA-seq.

**Figure 3.**
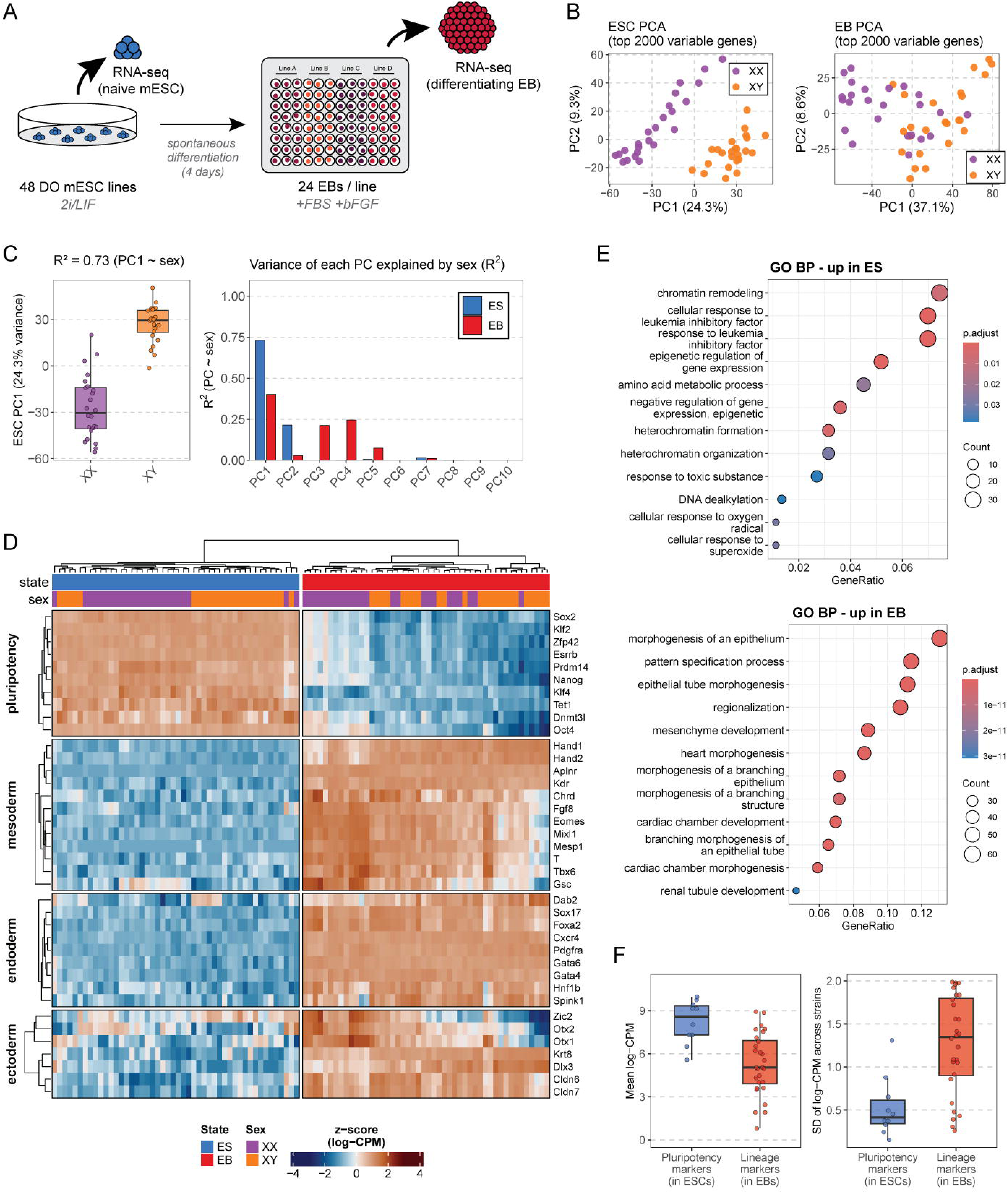
Characterizing variation in pluripotency and differentiation propensity in the DO mESC panel. (A) Overview of the DO mESC to EB differentiation protocol. RNA was extracted from cells in the pluripotent state (ESC) and again in the differentiating state (EB) after 4 days of growth for differential gene expression analysis. (B) First two principal components of gene expression (RNA-seq) from the top 2000 variable genes in DO ESCs (left panel) show the primary source of transcript variability is chromosomal sex (24.3%). First two principal components of gene expression from the top 2000 variable genes in DO EB cells (right panel). (C) PC1 variance is strongly attributable to chromosomal sex in the ESC state (R^2^ = 0.73) but much weaker in the EB state (R^2^ = 0.40) (left panel). Variance of the first 10 PCs explained by sex between cell states (right panel). (D) Heatmap of transcript expression for individual marker genes (rows) in log-CPM followed by z-score normalization centered on per-gene mean for all 96 samples (ESCs and EBs); columns are clustered. Marker genes selected span pluripotency, the primed-pluripotency transition, and the three germ layers. (E) Gene ontology (GO) enrichment of biological processes (FDR-adjusted *P*-value < 0.05) shows an over-representation of pluripotency-associated signatures in the ESC state, while the EB state is enriched for pattern specification, regionalization, and various early heart-morphogenesis signatures. The color scale represents the FDR-adjusted *P*-value, with different magnitudes for each cell state. (F) Analysis of strain variability between pluripotency markers in ESCs and lineage markers in EBs. SD of log-CPM across strains in each state shows low inter-strain variability for pluripotency markers in ESCs (median SD = 0.38 log-CPM) compared to elevated variability for lineage markers in the EBs (median SD = 1.38) (right panel). Mean log-CPM for each marker set within its respective state is shown for comparison (left panel).

Principal component analysis of the 48 DO mESC lines profiled in both states revealed differences in transcriptional variation between pluripotent and differentiated cells. In ESCs, PC1 (24.3% variance) was strongly associated with chromosomal sex (R² = 0.73), establishing sex as the principal organizing axis of the naive ESC transcriptome under 2i/LIF conditions (Figure 3B–C). Following spontaneous EB differentiation, PC1 captured a comparable fraction of total variance (37.1%) but exhibited a weaker association with sex (R² = 0.40), with sex-attributable variance redistributed across lower-ranking components (e.g., PC3 and PC4, R² = 0.21 and 0.25, respectively). Aggregated across the top ten principal components, sex accounted for approximately 20% of total transcriptional variance in ESCs and 17.5% in EBs (Supplemental Table 4). Repeating the analysis separately for autosomal and sex-linked genes showed the attenuation was specific to the sex chromosomes (57.0% in ESCs versus 44.7% in EBs; 21.6% relative reduction), with autosomal sex effects essentially unchanged between states (13.1% versus 14.4%; Supplemental Table 5).

We next examined expression of 38 curated marker genes spanning naive pluripotency, the primed-pluripotency transition, and the three germ layers across the 48 DO mESC panel lines in both ESC and EB states (Figure 3D). In the naive ESC state, all pluripotency markers, including the canonical core (*Oct4*, *Sox2*, *Nanog*), naive-specific transcription factors (*Klf2*, *Esrrb*, *Prdm14*, *Zfp42*), and chromatin regulators (*Tet1*, *Dnmt3l*), were uniformly and highly expressed across all 48 strains, consistent with maintenance of the naive pluripotency network under 2i/LIF conditions (Figure 3E, top panel)^43–46^. Following four days of EB differentiation, these markers were sharply downregulated (*Esrrb* log₂FC = −6.50; *Zfp42* log₂FC = −6.50; *Klf2* log₂FC = −5.69; all FDR < 0.001), with induction of the post-implantation epiblast marker *Fgf5* (log₂FC = 5.96; FDR < 0.001), indicating exit from naive pluripotency and differentiation toward a primed/formative state^47^.

Markers of all three germ layers^46^ were analyzed in EBs. Mesoderm specification was reflected by primitive-streak markers (*T, Mixl1, Eomes, Fgf8, Gsc, Chrd*; log₂FC range 3.02–7.79; FDR < 0.001) and an emerging cardiac/extraembryonic mesoderm program (*Hand1* log₂FC = 10.70; *Hand2* log₂FC = 8.33; *Aplnr* log₂FC = 13.24; *Kdr* log₂FC = 6.95), the latter consistent with the heart-morphogenesis and endocardial signatures observed in GO enrichment analysis of EB-upregulated genes (Figure 3E, bottom panel). Endoderm formation was supported by definitive-endoderm markers (*Sox17* log₂FC = 7.55; *Cxcr4* log₂FC = 7.30) alongside primitive/visceral-endoderm markers (*Gata4, Gata6, Dab2, Spink1*; log₂FC range 3.06–6.75). Ectoderm induction, marked by *Otx2, Dlx3,* and *Nes*, showed lower magnitude. Together, these data confirm coordinated pluripotency exit and multilineage germ-layer specification across the DO mESC panel.

To assess genetic variability in pluripotency maintenance and lineage induction, we compared expression variation between lines for pluripotency markers in ESCs and germ-layer markers in EBs. Pluripotency markers displayed uniformly low variation across lines (median SD = 0.38 log-CPM), whereas germ-layer markers in EBs varied substantially (median SD = 1.38 log-CPM; Figure 3F, right panel). Several primitive-streak and cardiac mesoderm regulators (*T, Mixl1, Eomes, Hand1*) exceeded SDs 1.8, equivalent to ∼3.5-fold expression difference between lines. While lineage markers were expressed at lower mean levels within EBs than pluripotency markers in ESCs (Figure 3F, left panel), this difference is consistent with their restriction to heterogeneous subpopulations within EBs. The elevated inter-line variability of these markers therefore suggests that genetic background influences the magnitude, cellular proportion, or timing of lineage induction.

### Genetic variation shapes the transcriptional response to cadmium exposure

To demonstrate the utility of this panel for genetic mapping, we tested how genetic background shapes the transcriptional response to an environmental exposure. Cadmium (Cd) is a toxic heavy metal that negatively affects gestational and birth outcomes, dysregulates organs, and interferes with early cell fate decisions, neurulation, and neuronal cell populations in humans and mouse^30,48–51^. However, many previous *in vitro* toxicogenomic experiments relied on single, isogenic lines with limited ability to elucidate complex regulatory networks, leaving the teratogenic potential of Cd in diverse backgrounds and early development incompletely understood^30^.

Cd exposure of DO founder EBs revealed that 75.0% of the variation in EB size is attributable to genetic background (Figure 4A–B), indicating substantial heritable variation in Cd response and establishing EB area as a tractable readout for concentration selection. We therefore conducted concentration-response modeling on a subset of lines (Figure 4C). Seven concentrations, ranging from 1 µM to 40 µM, were tested on nine DO mESC lines throughout our previously described 4-day EB differentiation protocol. The change in EB area in Cd-exposed versus unexposed control lines, acquired from segmentation on up to 48 EBs per line, was used as a readout for Cd toxicity (Figure 4D– E). Two independent experiments determined an effective concentration producing 20% response (EC20) of 4.4 µM (Figure 4F) which was used for the Cd toxicogenomic screen. To determine the genetic basis of Cd-induced transcriptomic changes, we differentiated 200 DO mESC lines into EBs in the presence or absence of 4.4 µM Cd and performed bulk RNA-seq. Following quality control checks, all 200 lines were retained for mapping and haplotype reconstruction. Because each cell line contributes control and exposed samples, this paired design permits mapping of steady-state expression within each condition as well as of the change in expression between them (i.e. the response to Cd exposure). We therefore performed eQTL mapping on the control and Cd-exposed conditions separately, and additionally mapped gene-by-environment (G×E) eQTL after adjusting for control expression. We mapped 7007 (control), 7152 (exposed), and 1916 (G×E) significant eQTL (autosomal LOD > 8.2, X-chromosome LOD > 7.87) influencing 6533, 6710, and 1836 genes, respectively. The smaller number of G×E eQTL is expected as any variant altering expression similarly in both conditions is removed by adjustment for control expression, so the loci that remain are those whose effects depend on Cd exposure.

**Figure 4.**
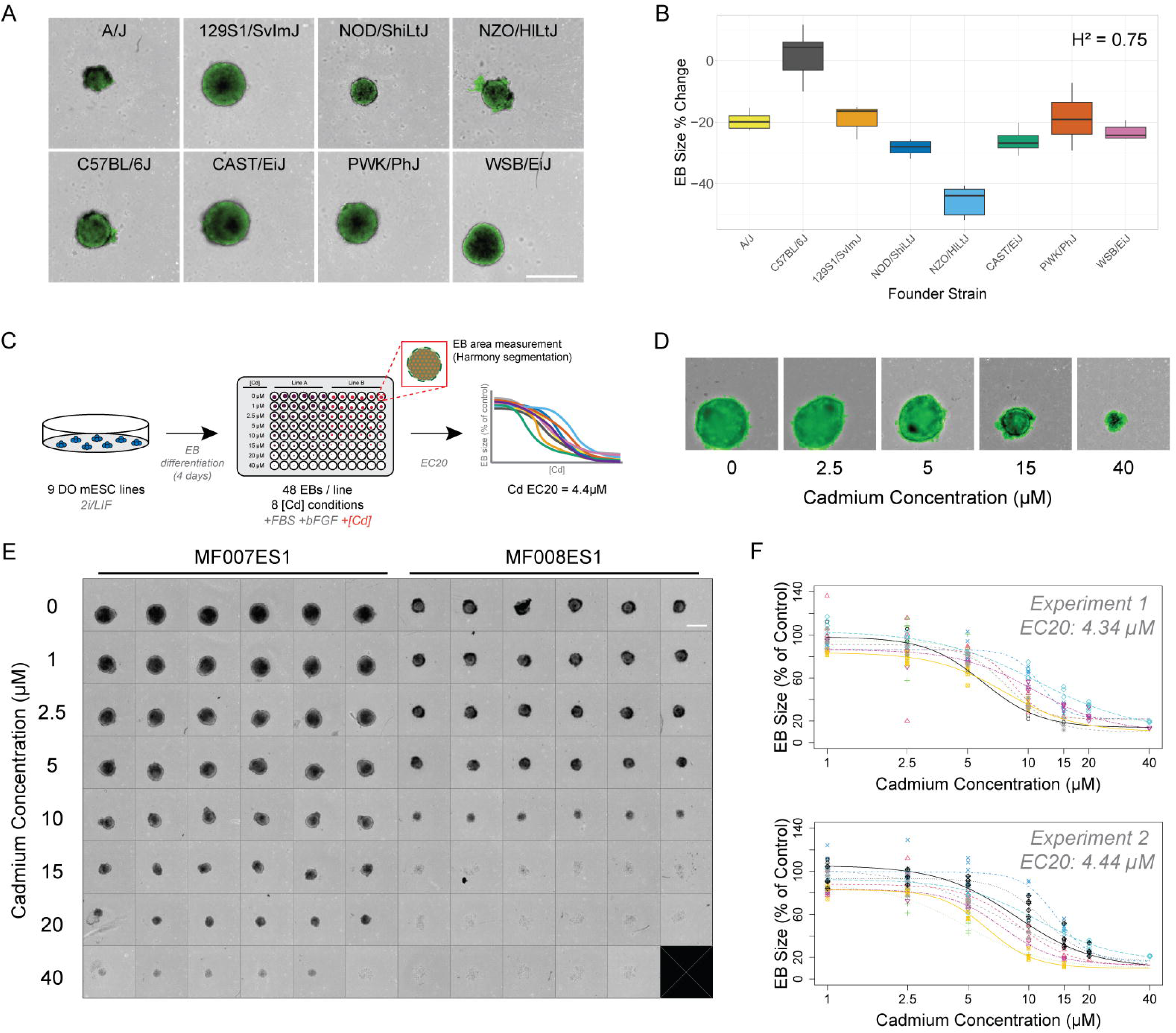
DO founder, mESC line EB segmentation and concentration response modeling. (A) Representative images from segmentation of all 8 DO founder strains following 4 days of differentiation to EBs. Identified EB objects are highlighted in green, representing the area measured for analysis. Scale bar set to 500 µm. (B) Plot showing percent change in EB size from all 8 founder strains after dosing with 4.4 µM Cd (percentage derived from mean exposed – mean control). The calculated heritability score indicates 75% of the variation in percent change in EB size seen between founders after Cd dosing is due to differences in genetic background. (C) Overview of the EB Cd concentration response modeling protocol. 48 EBs were differentiated for 4 days from 9 DO mESC lines under 7 different Cd concentrations plus a control. The change in EB area in exposed versus control lines was measured and used as a proxy for Cd toxicity. (D) Segmentation of selected EBs grown under different Cd concentrations. EB region highlighted in green. (E) Representative image of a 96-well plate containing EBs from 2 different DO mESC lines at different Cd concentrations. Black squares indicate failed imaging and well contents were excluded from analysis. Scale bar set to 500 µm. (F) Concentration response modeling curves from 2 separate experiments used to calculate the final EC20 of 4.4 µM for the Cd toxicity screen.

In the eQTL map, each point denotes a significant QTL positioned by the genomic location of the QTL on the x-axis and of its target gene on the y-axis; points falling along the diagonal therefore represent genes regulated by variants near their own locus. Approximately 75% of significant eQTL were in proximity (within ±2 Mb) to their target gene and are classified as local eQTL in both control (5277/7007; 75.3%) and exposed (5394/7152; 75.4%) conditions. By contrast, only 30.6% (587/1916) of significant G×E eQTL were local (Figure 5A, Supplemental Figure 2). This asymmetry is consistent with established differences between local and distant regulatory variation. Local effects are typically condition-invariant, detectable across tissues and cell states, whereas distant effects more frequently depend on cellular context^53^. Most local eQTL are therefore expected to act similarly in control and exposed EBs and to be removed by adjusting for control expression, despite their large effect sizes. Notably, the local G×E eQTL that persists represents a class of variant whose effect on its own gene is responsive to Cd exposure.

**Figure 5.**
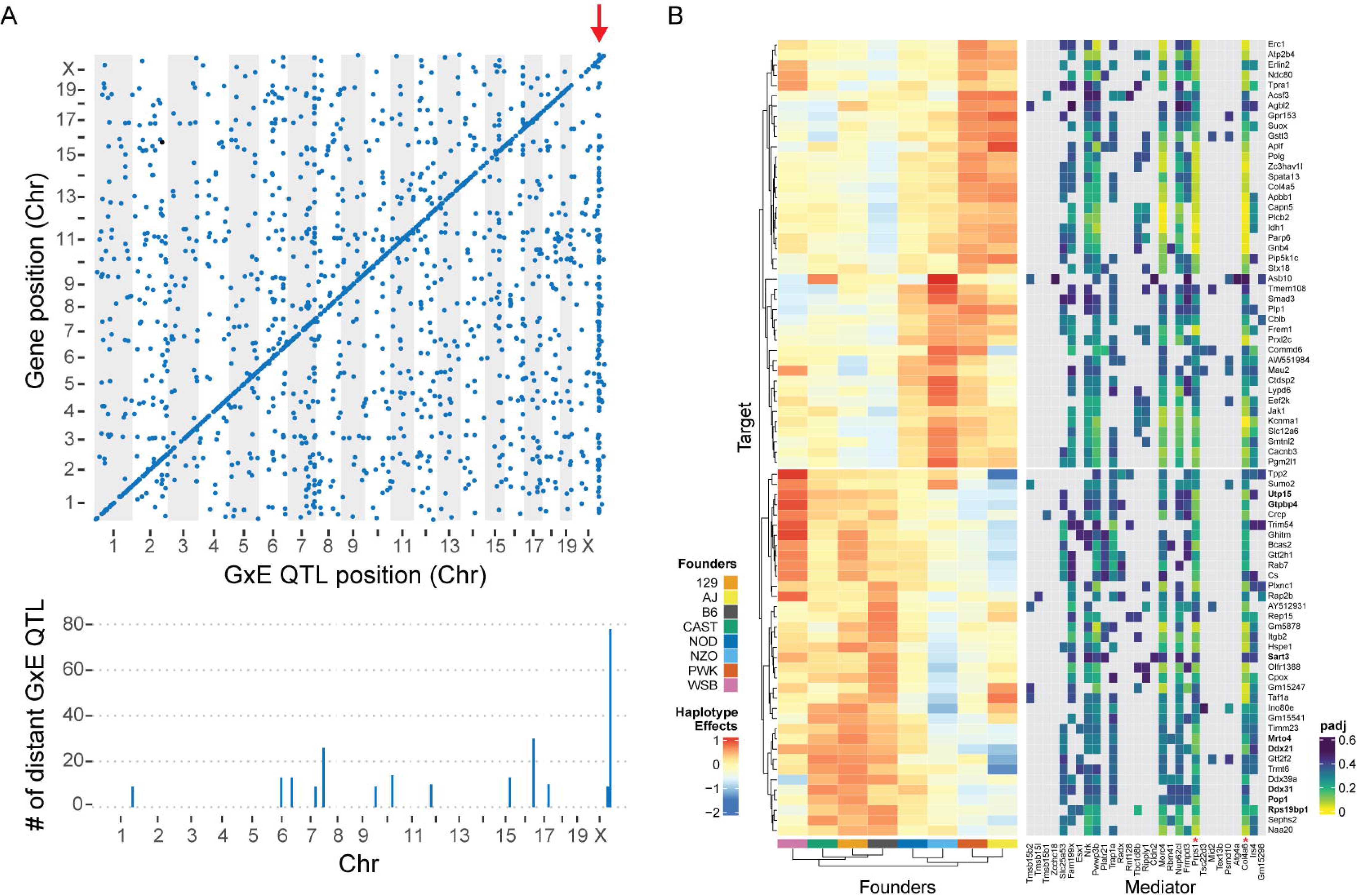
Validating the DO mESC panel as a QTL mapping and toxicogenomic resource. (A) Scatterplot of genomic locations of individual G×E QTL (x-axis, autosome LOD > 8.2, X chromosome LOD > 7.87) versus location of target genes (y-axis) for EBs. 1916 significant QTL were mapped influencing 1836 target genes. Genetic effects acting locally fall along the diagonal line, while effects acting distally lie off the diagonal. Most significant G×E QTL (69.4%) map far from their target gene. 13 G×E QTL “hotspots” were mapped (bottom panel), with the largest hotspot appearing as a vertical line on the X chromosome (red arrow) and influencing 78 target genes (top panel). (B) Founder haplotype effects for the X chromosome hotspot show two clusters with variation driven by different founders (left panel). Red indicates higher target gene expression in EBs carrying a haplotype derived from one of eight DO founder lines; blue indicates lower target gene expression. Mediation analysis identified two candidates, *Prps1* and *Col4a6* (red asterisks). ORA-identified target genes associated with ribosomal biogenesis and rRNA processing pathways shown in bold (right panel).

We next asked whether distant G×E eQTL co-localized to discrete genomic loci, which would indicate that variation at a single locus shapes the Cd response of many genes. Such eQTL “hotspots” appear as vertical bands in the eQTL map; we identified 13 G×E eQTL hotspots, including a major hotspot on the X chromosome that accounts for expression variation in response to Cd exposure for 78 target genes (Figure 5A). To determine if the affected genes share a common biological function, we performed overrepresentation analysis (ORA) on the suggestive target gene sets (*n* = 257) of the X chromosome hotspot (Supplemental Table 6), which identified enrichment for pathways involved in ribosomal biogenesis and rRNA processing. Because each DO genome is a mosaic of the eight founder haplotypes, the effect of each founder allele at an eQTL can be estimated separately for every target gene. Clustering the founder haplotype effects for each of the 78 significant target genes revealed two clusters driven by different sets of founders (Figure 5B). In one cluster, target gene expression was elevated in EBs carrying WSB/EiJ, CAST/EiJ, 129S1/SvImJ, and C57BL/6J haplotypes at the X chromosome hotspot. This cluster contains the target genes driving the ribosome biogenesis and rRNA processing enrichment. Together, these results indicate that a single X-linked locus modulates the Cd response of a coherent set of ribosome biogenesis genes, in a manner dependent on founder haplotype.

A distant eQTL is expected to act through a gene near the QTL itself, whose own expression or function varies with genotype and in turn influences expression of the distant targets. To nominate such intermediaries, we tested whether expression of each gene local to the hotspot statistically accounts for (i.e., mediates) the variation in each target gene expression. Two genes were identified as candidate mediators, *Prps1* and *Col4a6* (Figure 5B). *Prps1* was the strongest mediator for the target genes in the ribosome biogenesis pathways identified by ORA. Notably, *Prps1* itself has a local eQTL detected in the G×E analysis, indicating that the effect of genotype on *Prps1* expression was after Cd exposure. Together, these analyses demonstrate that the DO mESC panel can resolve environmentally responsive regulatory variation, coordinated target gene sets, and candidate mediators, providing a foundation for mechanistic follow up.

## Discussion

Expanding the diversity of cellular systems genetics resources is essential for identifying genes and regulatory networks contributing to complex traits and ultimately generating insights that are broadly relevant across genetically diverse populations. Here, we generated and characterized a panel of 299 genetically diverse mESC lines obtained from unique, non-sibling, highly outcrossed DO population mice. To maximize the capture of genetic background effects on phenotypes of interest, all lines can be grown in identical, defined, feeder-free and serum-free 2i/LIF culture conditions. Using the high-density GigaMUGA SNP array, we confirmed the highly heterozygous, outbred genomes of the DO population are well represented across the panel, with minimal kinship and balanced chromosomal sex.

Within the ESC state, we found chromosomal sex to be the dominant source of transcriptional variation, and this association attenuated following differentiation to EBs. This is consistent with initiation of X-inactivation during post-implantation development^54^ and aligns with previous observations that XX mESCs do not differentiate until X-inactivation has occurred, resulting in a delayed exit from pluripotency relative to XY lines^55,56^. Indeed, across many mammalian species, XX embryos develop more slowly than age-matched XY embryos^57,58^. Within EBs, a subset of XX lines with reduced expression of the cardiovascular mesoderm markers *Hand1* and *Hand2* showed correspondingly higher expression of pluripotency and primitive-streak genes, implying that certain genetic backgrounds are biased toward an earlier developmental state. In this way, chromosomal sex may contribute to differences in developmental progression in DO mESC lines, with XX EBs predominantly occupying an earlier gastrulation-stage state while XY EBs more closely resemble later gastrulation and early neurulation-stage progenitors. However, even among lines of the same chromosomal sex, DO EBs displayed substantial transcriptional variation.

We found that lineage marker expression in EBs varied substantially more between lines than pluripotency marker expression in ESCs, implying that genetic background exerts greater influence on differentiation than on naïve pluripotency. Specifically, expression of the lateral plate and cardiovascular mesoderm markers *Aplnr* and *Kdr* was consistently high across lines, whereas earlier primitive-streak markers (e.g., *T, Mixl1, Eomes, Fgf8, Gsc,* and *Chrd*) showed variable expression. Previous single-cell profiling studies have reported that differentiated EBs contain overlapping developmental stages and retain subpopulations of pluripotent-like or epiblast-like cells, even as mesodermal lineages emerge^59,60^. Further, inbred founder strains of the DO differ in gestational timing and lineage commitment^61,62^, suggesting the panel reproduces developmental variability seen in the DO source population. Distinguishing the underlying basis of this variability requires resolving whether elevated strain-level variability reflects differences in the proportion of cells adopting each lineage or differences in the rate at which lines progress along a developmental trajectory, a distinction inaccessible to bulk measurements.

While genotypes and gene expression data for a subset of the untreated EBs here have previously been used in a study benchmarking genotyping methods for haplotype reconstruction in complex mouse crosses^63^, this present work demonstrates the power of the DO mESC panel for identifying interactions between genotype and environment. We performed G×E eQTL mapping of Cd exposure during EB differentiation and identified 13 trans-regulatory hotspots, the largest of which on chromosome X regulated 78 genes. The target genes of this hotspot were enriched for ribosome biogenesis, ribonucleoprotein complex biogenesis, and rRNA processing pathways. Ribosome biogenesis and rRNA processing are core nucleolar functions that are disrupted during nucleolar stress^64–67^. Given that Cd exposure induces oxidative stress and DNA damage response pathways^68–70^, this enrichment suggests that genotype-dependent differences in gene expression in response to Cd exposure may modulate nucleolar regulatory programs during differentiation.

Mediation analysis identified two candidates, *Prps1* and *Col4a6,* as potential mediators of this X chromosome eQTL hotspot. *Prps1* catalyzes the synthesis of phosphoribosyl pyrophosphate (PRPP), an essential precursor for nucleotide synthesis^71,72^, while *Col4a6* encodes a component of type IV collagen, a key constituent of extracellular matrix (ECM) basement membranes and connective tissue^73,74^. Of these, *Prps1* function aligns with the enriched pathways because rRNA synthesis consumes ribonucleotides at high rates during ribosome assembly. A role for Cd in perturbing these pathways is supported by studies in mouse embryos and human cells showing that Cd and other heavy metal toxicants disrupt ribosome biogenesis, including pre-rRNA processing and rRNA synthesis^48,75^.

The link observed here between Cd exposure, *Prps1*, and rRNA processing is notable given the importance of nucleotide homeostasis during early embryonic development. The peri- and post-implantation stages are characterized by extensive cellular proliferation and biosynthesis, placing increased demands on nucleotide availability and ribosome production. Disruption of either nucleotide homeostasis or ribosome biogenesis can impair developmental progression and lead to embryonic lethality^76–81^. Redox signaling in the early embryo is tightly regulated, with reactive oxygen species (ROS) contributing to normal control of differentiation and proliferation^70,82^. Cd- induced ROS may disturb this balance, damaging nucleic acids and increasing demand for nucleotides for repair and replacement^83^. Given the high nucleotide requirements of rRNA transcription and ribosome assembly, variation in *Prps1* activity under Cd-induced oxidative stress could influence the ability of differentiating cells to sustain nucleolar functions. While this mechanism remains speculative, the combination of pathway enrichment and mediator candidates supports a role for one or more regulatory pathways in shaping the Cd response during early embryonic differentiation.

Beyond the specific findings reported here, a primary contribution of this work is the resource itself. Existing DO mESC lines are privately held and were derived from fewer DO founders at earlier outbreeding generations, resulting in substantial kinship that can reduce mapping power^4,32^. By deriving each line from a unique, non-sibling mating, we minimized relatedness across the panel while capturing the allelic diversity of the population. Critically, each DO mouse is a genetically unique individual whose combination of alleles cannot be recovered once the animal is gone; however, pluripotent cell lines convert these into a renewable resource. Genotyping is therefore performed once here as part of resource development, enabling cellular phenotypes to be measured and accrued to the same set of genomes. Investigators need only to assay a phenotype of interest and proceed directly to mapping without animal husbandry, genotyping infrastructure, or regulatory burden that *in vivo* studies require. While cell-based systems access a narrower range of phenotypes than living animals, loci identified *in vitro* can be followed up in mice to functional validation, an option unavailable in human pluripotent cell panels.

Collectively, these results establish the DO mESC panel as a powerful resource for developmental systems genetics. All DO mESC lines reported here are available through The Mutant Mouse Resource & Research Centers (MMRRC) Repository under the JAX Diversity Outbred mESC collection. This sex-balanced panel maintains the genetic heterozygosity found in the DO mouse population and recapitulates key features of early development, including expected naïve pluripotency and lineage marker expression, and known sex-specific dynamics during early cell-state transitions. Its genetic diversity supports high-resolution mapping of loci, regulatory networks, and candidate mediators underlying developmental and genotype-by-environment phenotypes, and the toxicogenomic approach demonstrated here with cadmium extends readily to other exposures, differentiation protocols, and cell states. Ultimately, this panel provides a powerful framework for understanding how combinations of genetic variants shape susceptibility to environmentally influenced traits and complex diseases.

## Resource Availability

### Lead contact

Requests for further information and resources should be directed to the corresponding author and lead contact, Christopher L. Baker.

### Materials Availability

All cell lines are available through The Mutant Mouse Resource & Research Centers (MMRRC) Repository (https://www.mmrrc.org) under the “JAX Diversity Outbred mESC” collection.

### Data and Code Availability

RNA-seq data from panel mESCs and differentiated EBs used in all experiments are available through NCBI Gene Expression Omnibus (accession number: GSE344387). QTL mapping objects are available through Figshare (10.6084/m9.figshare.33546949).

## Acknowledgements and Funding

We gratefully acknowledge Genome Technologies at the Jackson Laboratory for their expert RNA-seq library preparation and sequencing services that supported this study. Scientific services at the Jackson Laboratory are supported in part through the National Cancer Institute’s Cancer Core Grant P30CA034196. We would also like to thank Melissa L. Berry and the Reproductive Sciences staff at the Jackson Laboratory for their dedicated panel curation and cryopreservation efforts. In addition, we would also like to thank the members of the Reinholdt, Baker, and Munger laboratories for their guidance, input, and support.

Generation of this resource was supported by the National Institutes of Health Office of Research Infrastructure Programs under award number R24OD030037. MMB and CLB were supported by the National Institute of General Medical Sciences under award number R35GM133724. The content is solely the responsibility of the authors and does not necessarily represent the official views of the National Institutes of Health.

## Author Contributions

Conceptualization: LGR, SCM, CLB. Data curation: MMB, ES, KJ, SJW, HBD, SA, WM, AS, LGR, SCM, CLB. Formal analysis: MMB, ES, SJW, HBD, SA, AS, LGR, SCM, CLB.

Funding acquisition: LGR, SCM, CLB. Investigation: ES, KJ, WM, CB. Methodology: ES. Project administration: MMB, ES. Resources: MMB, ES, KJ, WM. Software: HBD. Supervision: DMG, LGR, SCM, CLB. Validation: MMB, ES, SJW, HBD, SA. Visualization: MMB, ES. Writing – original draft: MMB. Writing – review & editing: MMB, LGR, SCM, CLB.

## Methods

### Panel derivation

DO mice (RRID:IMSR_JAX:009376, JAX stock 009376) were obtained from the Jackson Laboratory. All animal experiments were approved by the Animal Care and Use Committee at the Jackson Laboratory (Animal Use Summary #20030). DO mESCs were derived as described previously^31^ with modifications. Briefly, DO mESCs were derived by breeding non-sibling male and female DO mice, aged approximately 6 weeks, and culturing embryonic day 3.5 (E3.5) blastocysts. Blastocysts were initially collected in M2 culture media (Gibco) before being pipetted onto plates containing 0.1% gelatin (Stem Cell Technologies) and irradiated MEF feeders in 2i/LIF/low-serum ES media, comprised of a 50% DMEM/F12 (Gibco) and 50% Neurobasal medium (Gibco) base supplemented with 0.5X N-2 (Gibco), 0.5X B27 (Gibco), 1X PenStrep (Gibco), 0.05% BSA (Gibco), 2 mM GlutaMAX (Gibco), 150 µM monothioglycerol (Sigma), 1000 U/mL LIF (R&D Systems), 1 µM PD0325901 (Stem Cell Technologies), 3 µM CHIR99021 (Stem Cell Technologies), and 2% ES-grade FBS (Gibco). Following outgrowth formation (∼5–10 days after plating), cells were disaggregated and passaged into a 12-well plate treated with 0.1% gelatin (Stem Cell Technologies) and irradiated MEF feeders in 2i/LIF/low-serum ES media. Cultures were weaned off MEF feeders during derivation and established at P4–6, then frozen in 50% culture media, 40% FBS, and 10% dimethyl sulfoxide (DMSO). All mESC sample cryovials were transferred for long-term storage to liquid nitrogen holding tanks after controlled freeze (-1°C/min) for 24–48 hours at -80°C. Lines were further expanded from frozen stocks for genotyping, contaminant screening, and biobanking. Derivation and cell cultures were grown at 37°C. All lines tested negative for bacterial and fungal growth and for the presence of mycoplasma using a PCR detection system. Chromosomal sex of all 299 derived cell lines can be found in Supplemental Table 1.

### ESC culture and differentiation to embryoid bodies

*ESC culture:* Undifferentiated mESC lines were thawed at P6–12 and cultured for 2 days in the 2i/LIF ES media conditions described above, but without the inclusion of serum (2i/LIF/serum-free). After 48 hours, cells were either differentiated to EBs or snap frozen in liquid nitrogen and stored at -80°C prior to RNA isolation.

*EB differentiation in the presence or absence of Cd:* ES cells were dissociated with TrypLE (Gibco), then counted. The ASSIST Plus pipetting robot system (INTEGRA Biosciences) was used to seed cells into 96-well spheroid ultra-low attachment plates (Perkin Elmer) in embryoid body medium at 750 cells/well. Embryoid body medium consisted of a DMEM 4.5 g/L (Gibco) base supplemented with 15% FBS, 2 mM GlutaMAX-1 (Gibco), 1X PenStrep (Gibco), 1% NEAA, 1% sodium pyruvate, 0.1 mM 2- mercaptoethanol and 12 ng/mL bFGF in the presence or absence of 4.4 µM cadmium starting at the time cells were seeded (day 0 of differentiation). Plates were spun at 1200 RPM for 3 minutes to aggregate cells. EB cultures were grown for 4 days at 37°C, with a half media change on day 2. For each cell line, up to 24 EBs (1 EB/well) were pooled per condition (unexposed vs. exposed) and snap frozen in liquid nitrogen and stored at -80°C prior to RNA isolation.

### Genotyping and genetic characterization

#### Genotyping

DNA was collected using the DNeasy Blood and Tissue Kit (Qiagen) from each DO mESC line at expansion. Genotyping was performed by Neogen Genomics using the Giga Mouse Universal Genotyping Array (GigaMUGA)^29^ on the Illumina Infinium platform. A total of 111,092 SNP markers passed quality control and were used for haplotype reconstruction and eQTL mapping. Each of the 36 possible founder diploid genotype probabilities were estimated at each marker location using a hidden Markov model implemented in `r/qtl2`^84^, and 36-state diploid genotypes were condensed to 8 founder allele probabilities for eQTL mapping (see “eQTL Mapping” below).

### Founder strain representation

Genome-wide and chromosome-specific founder allele frequencies were estimated from inferred founder haplotype probabilities. For each founder strain, observed allele frequencies were compared with the expected 0.125 frequency using Chi-squared tests and resulting p-values were corrected for multiple testing using the Benjamini-Hochberg procedure. The relative contribution of each of the eight DO founder strains was summarized across autosomes and the X chromosome for all DO mESC lines.

### Kinship analysis

Genetic relatedness among DO mESC lines was assessed using a kinship matrix calculated from inferred founder genotype probabilities for autosomal markers using the calc_kinship() function in `R/qtl2`^85^. Kinship was estimated as the proportion of shared founder alleles between DO mESC lines. Sex chromosome markers were excluded from the kinship analysis.

### Sex inference

Sex was inferred from mean GigaMUGA probe intensities on the X and Y chromosomes. Samples were classified based on the relative X-chromosome dosage and the presence or absence of the Y-chromosome signal.

### Aneuploidy analysis

Autosomal aneuploidy analysis was carried out as previously described^36^, with the exception that a chromosome was only flagged as aneuploid if it was both a statistical outlier (|robust z| ≥ 4; outlier strength relative to the panel) and displayed an absolute chromosome-wide expression shift of ∼11% (0.15 log2FC vs. panel). Genome-wide B-allele frequency (BAF) and log R ratio (LRR) profiles were generated from GigaMUGA SNP array data and inspected for chromosome copy number changes. In diploid regions, SNPs are expected to cluster at BAF values of 0, 0.5, and 1. Regions exhibiting altered BAF distributions together with corresponding shifts in LRR signal intensity were interpreted as evidence of copy number variation or aneuploidy. Karyotypic analysis was performed on 20 DO mESC lines by Cell Line Genetics.

### Transcriptomic profiling for pluripotency and differentiation characterization

#### RNA extraction, library preparation, and sequencing

RNA was extracted according to manufacturer protocol using a NucleoMag RNA Kit (Macherey Nagel) and purified using a KingFisher Flex system (ThermoFisher). Library preparation was enriched for polyA containing mRNA by JAX Genome Technologies using the KAPA mRNA HyperPrep Kit (Roche Sequencing and Life Science). Paired-end bulk RNA sequencing was performed by JAX Genome Technologies on an Illumina NovaSeq 6000 with a 150 bp read-length and mean read depth of 40M reads per sample.

#### RNA-sequencing data processing, filtering, and normalization

Gene expression counts from GBRS (see “eQTL mapping” below) were combined for 48 DO mESC lines and their matched EBs (not exposed to Cd, as described above), yielding 32,176 genes across 96 samples. RNA-seq data were analyzed using the `R/edgeR` package (version 4.2.0). Lowly expressed genes were removed prior to downstream analyses. Library sizes were normalized using the trimmed mean of M-values (TMM) method, and expression values were transformed to log_2_ counts per million (logCPM) for visualization and variance analyses. Batch was excluded as a covariate because it was fully confounded with cell state. Differential expression between ESC and EB states was tested using edgeR quasi-likelihood. The design included chromosomal sex as an additive covariate (∼ sex + condition). P-values were adjusted by the Benjamini-Hochberg method, and genes were considered differentially expressed at FDR < 0.05 and |log_2_FC| > 1, where positive log_2_FC indicates higher expression in EBs.

### Principal component analysis

PCA was performed on within-state logCPM matrices using the 2,000 most variable genes by row variance. For each of the first ten components, the proportion of variance attributable to chromosomal sex was estimated as the R^2^ from a linear model of the component score on sex, then multiplied by that component’s percentage of total variance and summed across components. To assess whether sex effects arose from the sex chromosomes or the autosomes, the same procedure was repeated on matrices restricted to autosomal or sex-linked genes assigned by Ensembl annotation. Because fewer than 2,000 sex-linked genes were retained, all sex-linked genes were used in the PCA (n = 520 in ESCs, n = 533 in EBs), whereas the autosomal analyses used the 2,000 most variable genes per cell state.

### Gene ontology enrichment

Enrichment of biological process terms was tested separately for genes up- and down-regulated in EBs using *enrichGO* (GO.db 3.19.1, GO release 2024-01-17) from `R/clusterProfiler` (version 4.12.6)^86^ with the org.Mm.eg.db (3.19.1) annotation. For each direction, the 500 most significant differentially expressed genes by FDR were used as input, with all 15,887 genes retained in the joint filtered dataset as the background set. P-values were adjusted by the Benjamini–Hochberg method with a q-value cutoff of 0.05.

### Concentration response and heritability modeling

### Image acquisition and segmentation

Brightfield mESCs images were captured on an Axio Observer phase contrast fluorescence microscope (ZEISS) using a 10X objective and a 0.35X C-mount reduction adapter. mESC image scale bars were reconstructed using a counting hemocytometer with a Neubauer grid on the original AXIO Observer configuration. Images were converted to 8-bit grayscale in ImageJ^87^. 96-well plates containing EBs were imaged using the Opera Phenix High-Content Imaging System (PerkinElmer) with a 10X objective. A z-plane was acquired from 10 fields per well, and the z-stack maximum projection was analyzed and processed using Harmony 4.9 software with PhenoLOGIC (PerkinElmer). Segmentation was performed using Harmony 4.9 software, and images with more or less than one EB per well were not analyzed.

### Broad-sense heritability modeling

Broad-sense heritability (H²) was estimated using the `R/lme4` package (version 1.1-35.3)^88^. For each founder line, the percent change in EB area following cadmium treatment was calculated relative to unexposed controls. A linear mixed-effects model was fit with founder strain, passage number, and experimental date included as random effects. Passage number and experimental date were included to account for batch-related variation. Heritability was calculated as the proportion of total variance attributable to founder strain.

### Cadmium concentration-response modeling

Concentration-response modeling was performed in R using the `R/drc` (version 3.0-1)^89^ package to determine EC20. For each DO mESC line, EB sizes were normalized by dividing the mean EB area at each cadmium concentration (1–40 µM) by the mean EB area at 0 µM cadmium and expressed as a percentage of the control. Normalized EB size was modeled as a function of cadmium concentration using a four-parameter log-logistic model (LL.4). The lower asymptote parameter was set to ≥10 to prevent biologically implausible minimum EB sizes, and the Box-Cox transformation parameter was set to 0. EC20 values were estimated from a pooled LL.4 model using the same constraints. The mean EC20 (4.4 µM cadmium) from 2 independent experiments was used for toxicology screening.

### eQTL mapping

#### RNA-sequencing and expression quantification

RNA was extracted from up to 24 pooled embryoid bodies derived from 200 DO embryonic stem cell lines with and without exposure to cadmium. RNA extraction, library preparation, and sequencing were performed as described earlier. Gene expression and GigaMUGA genotyping data from these untreated samples were also used previously in a study benchmarking DO genotyping methods^63^. Read pairs were aligned separately to an “8-way” transcriptome using `Bowtie`^90^ as implemented within GBRS^91^. This 8-way transcriptome contains founder strain-specific versions of all annotated transcripts that incorporate strain SNPs from the Mouse Genomes Project^92^ onto the reference genome (Genome GRCm39; Ensembl annotations v105; SNPs release 2112_v8). Read pairs with shared alignments were then passed to `EMASE`^93^ to quantify gene expression. `EMASE` employs an expectation-maximization algorithm to account for potential reference bias and resolve multi-mapping reads in gene expression quantification. Next, gene-level counts from `EMASE` were normalized by variance stabilization transformation using `R/DESeq2`^94^. To reduce the likelihood of spurious eQTL stemming from highly variable but lowly expressed genes, we filtered the variance-stabilized counts to gene counts that summed across all samples to at least 3,000 and had at least 100 unique normalized count values among all samples. A total of 17,405 genes passed filtering and were rank-Z transformed to approximate a normal distribution for QTL mapping. GBRS-derived genotype probabilities were compared with GigaMUGA-derived probabilities to assess concordance across samples. GigaMUGA-derived genotype probabilities were used for downstream analyses.

### Expression QTL mapping

Expression QTL were mapped on normalized, transformed gene-level expression values described above for DO EBs with and without exposure to cadmium. The 8-state founder allele probabilities were used to fit a linear mixed-effects model with sex and outbreeding generation as additive covariates and leave-one-chromosome-out kinship matrices to account for relatedness among individuals. When searching for gene-by-environment (G×E) QTL, we adjusted for heritable steady-state variation in gene expression by first regressing out the normalized control (unexposed) expression value out of the exposed expression value and then rank-Z transformed the resulting residuals. We performed eQTL mapping using this rank-Z transformed, control adjusted phenotype, according to the model

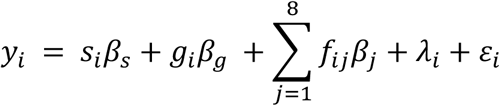

where *y_i_* is the rank-Z transformed gene expression value for <u>exposed</u> individual *i,* after adjustment for control expression, *β_s_* is the effect of sex, *s_i_* is the sex of individual *i*, *β_gg_* is the effect of outbreeding generation, *g_i_* is the outbreeding generation of individual *i*, *β_ii_* is the effect of founder allele *j*, *f_iii_* is the founder allele probability for founder *j* in individual *i*, and *λ_i_* is the kinship adjustment for individual *i.* This model was employed using the `R/qtl2` *scan1* function, and significant eQTL were detected using the *find_peaks* function when logarithm of the odds ratio (LOD) statistic exceeded a significance threshold of 8.2 for the autosomes and 7.87 for the X chromosome. LOD significance thresholds were determined by running *scan1perm* (with *perm_Xsp* set to true) on 750 randomly selected rankZ transformed control expression traits (1000 permutations each) and taking the median genome-wide threshold across these traits (significant 95%, suggestive 60%). Significant peak width was defined using the 95% Bayes credible interval from *find_peaks*. We assigned local eQTL as those with peaks within 2 Mb of the midpoint of the putatively regulated gene, and those outside of this range as distant eQTL.

### Gene-by-Environment QTL hotspots

We tallied all significant distant G×E QTL in windows of 50 markers with a 10-marker step size across each chromosome, assigning each peak to its nearest marker prior to counting, to identify loci where QTL were concentrated (or “hotspots”). Windows falling in the top 0.5% of significant peak density were then selected and collapsed into continuous bands to define final hotspot loci. To characterize the power of the panel to identify environmentally driven variation, we investigated one of the hotspots further. We identified the suggestive targets of the chosen hotspot and performed overrepresentation analysis (ORA) using `R/gprofiler2` ^95^ (version 0.2.4), and all expressed genes tested in the experiment as the background. For each target we extracted the founder allele effect coefficients using the `R/qtl2`^85^ (version 0.46) function ‘scan1blup’ and performed hierarchical clustering on the targets; then performed mediation analysis using the `R/intermediate`^96^ (version 0.2-4) function *mediation.scan* with candidate mediators defined as phenotypes local to the hotspot. QTLRetrievR^97^ (version 2.0.0.0) was used to integrate `qtl2` and `intermediate` for QTL mapping, haplotype effect estimates, and mediation analysis.

## Supplemental Information

**Supplemental Figure 1.**
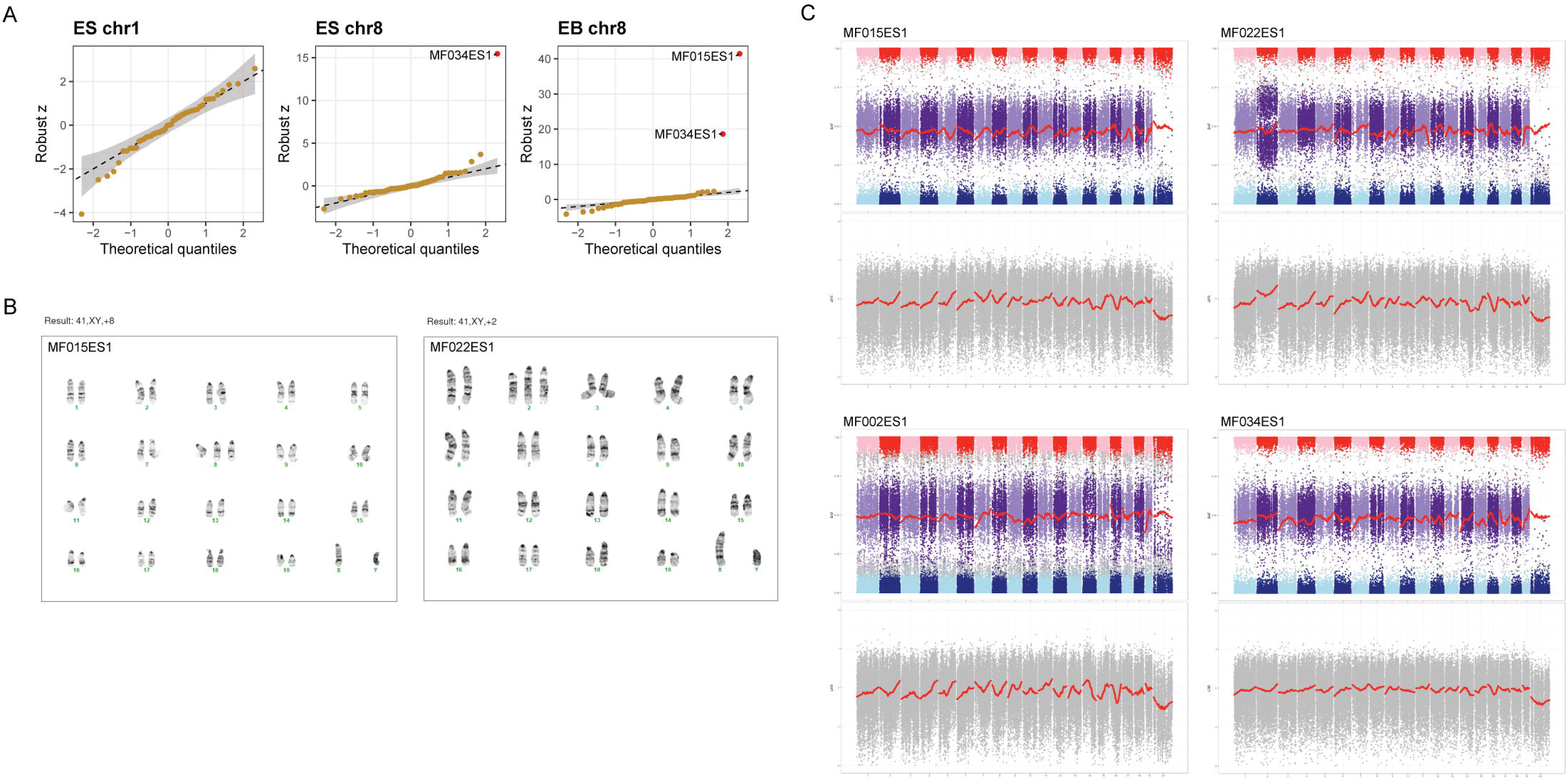
Additional aneuploidy analysis for select lines. (A) Quantile– quantile plots of the per-chromosome robust *z* against a normal reference, for a euploid chromosome (chr1, ESC) and the recurrently gained chr8 in each state using gene expression data. Points within the grey band match the expected null. (B) Metaphase spreads for MF015ES1 and MF022ES1 showing trisomy for chromosome 8 and 2, respectively. (C) B-allele frequency plots and Log R ratio (LRR) plots from GigaMUGA genotyping data confirm copy number increases in MF022ES1 (Chr 2) and MF002ES1 (Chr 14) but not MF015ES1 or MF034ES1.

**Supplemental Figure 2.**
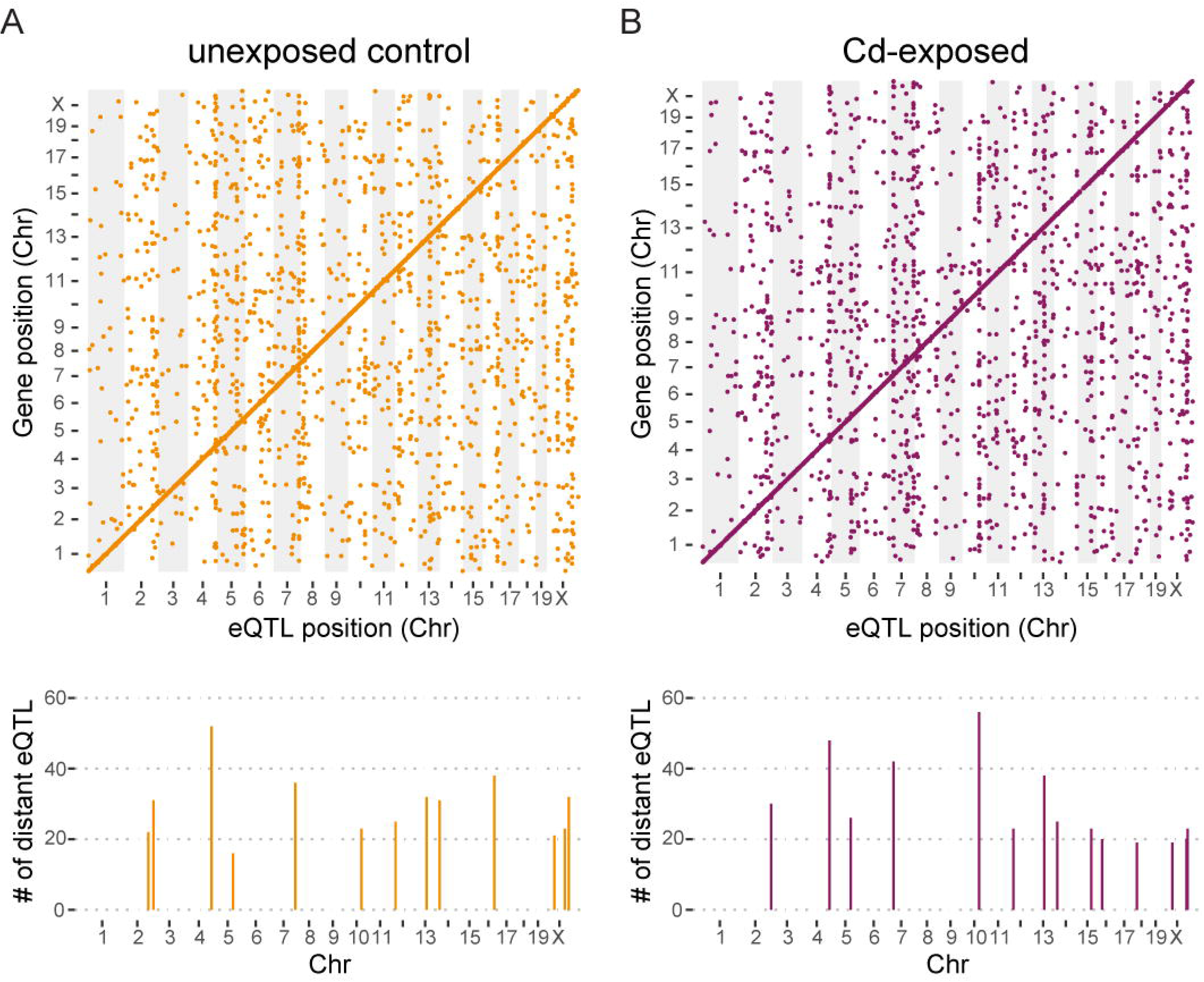
eQTL mapping on control and Cd exposed conditions. Scatterplot of genomic locations of control and exposed eQTL (x-axis, autosome LOD > 8.2, X chromosome LOD > 7.87) versus location of target genes (y-axis) for EBs. For the control condition, 7007 significant QTL were mapped influencing 6533 target genes, while the Cd exposed condition mapped 7152 significant QTL influencing 6710 target genes (top panels). eQTL hotspots for each and the number of target genes influenced shown below (bottom panel).

**Supplementary Table 1. Complete list of DO mESC panel lines and associated data.** Cell line naming convention lists the breeder pair (e.g., MF007 or MF111), followed by the embryo identifier (ES1 or ES2) from which the mESC line was derived. Strain of origin, chromosomal sex, cell passage, outcross generation, BSID#, GigaMUGA ID, MMRRC ID, and MMRRC strain name are included for each DO mESC line. Note that the GigaMUGA ID and BSID numbers do not always match the breeder pair ID (e.g., cell line MF012ES1 has a BSID# of CL9376x630 and a GigaMUGA ID of D0_003).

**Supplemental Table 2. Autosomal aneuploidy calls across both cell states.** Flagged whole-chromosome dosage events for autosomes only (off-diagonal points). All identified chromosomal changes are gains. Three gains were detected across both states, MF002ES1 (chr14), MF022ES1 (chr2), and MF034ES1 (chr8), consistent with a clonal aneuploidy carried through differentiation. One event is EB-specific where MF015ES1 gains chr8. Due to the number of lines analyzed (*n* = 48), a chromosome was flagged as aneuploid only if it is both a statistical outlier (|robust z| ≥ 4) and shifted by |chr_log2fc| ≥ 0.15 (∼11% expression shift), rather than using a rank-based *p*-value alone.

**Supplemental Table 3. Cytogenetic karyotyping results for 20 DO mESC lines**. For each line, 20 metaphases were counted, with 6/20 lines showing <70% euploid cells. 2/6 lines identified were XO.

**Supplemental Table 4. Cumulative percentage of transcriptional variance attributable to chromosomal sex across the top 10 principal components, by cell state.** The EB state showed less sex-attributable variance in the 10 summed PCs than the ES state (17.52% vs. 19.87%, respectively).

**Supplemental Table 5. Sex-attributable transcriptional variance in separate analyses of autosomal and sex-linked genes.** This variance was markedly higher on the sex chromosomes than on autosomes in both cell states. Values report the cumulative percentage of transcriptional variance attributable to chromosomal sex across the top 10 principal components, computed from compartment-specific PCAs (autosomal: chromosomes 1-19; sex chromosomes: X and Y). Change is reported in percentage points (pp) and relative to the ESC value.

**Supplemental Table 6. Overrepresentation analysis (ORA) of X chromosome hotspot suggestive and significant target genes.** Enrichment analysis was performed on 257 genes from the suggestive target gene set of the X chromosome hotspot. ORA identified significant enrichment of pathways involved in ribosome biogenesis, rRNA processing, and ribonucleoprotein complex assembly. The suggestive target gene set includes all 78 significant target genes.

